# *Cellulomonas zeae* sp. nov., *Lelliottia zeae* sp. nov., *Paraburkholderia zeae* sp. nov. and *Sphingomonas zeigerminis* sp. nov., four new bacterial species isolated from *Zea mays* and other hosts

**DOI:** 10.1101/2021.04.16.440235

**Authors:** Sarah Seaton, Jacqueline Lemaire, Patrik Inderbitzin, Victoria Knight-Connoni, Martha E. Trujillo

**Affiliations:** Indigo Ag, Inc., 500 Rutherford Avenue, Boston, MA 02129, United States; Departamento de Microbiología y Genética, Campus Miguel de Unamuno, University of Salamanca, Salamanca, Spain

## Abstract

Four novel bacterial species collected from healthy tissues of corn, rice and soybean plants in the United States are described. These include *Cellulomonas zeae* sp. nov. isolated from corn in Indiana; *Lelliottia zeae* sp. nov. from corn in Indiana and rice in Arkansas; *Paraburkholderia zeae* sp. nov. isolated from corn in Iowa; and *Sphingomonas zeigerminis* sp. nov. from corn in Mississippi and soy in Arkansas. No pathogenic strains are known for any of the novel species based on genome comparisons to assemblies in GenBank.

## INTRODUCTION

Bacterial communities present in soil account for the richest reservoir of biological diversity on Earth (Berendsen et al. 2012; Finkel et al. 2017). Many rhizosphere bacteria are of great importance due to their interactions with plants (Schenk et al. 2012), and this complex plant-associated microbial community is for the most part beneficial to the plant (Berendsen et al. 2012). Despite the importance of microorganisms for plants, these extremely complex microbial communities have remained largely uncharacterized (Schenk et al. 2012).

Maize or corn (*Zea mays* L.) is one of the most prevalent and economically important cereal crops worldwide used for human consumption, animal feed and as a raw material for energy production (Byrt et al. 2011). Maize is home to a rich diversity of plant-associated microorganisms that collectively account for roughly twenty times as many genes as the maize genome itself (Wallace and May 2018). Plant-associated microorganisms are increasingly being used for biotechnological applications, including biological control of plant pathogens, plant growth promotion, or isolation of active compounds (Ryan et al. 2008; Glick 2012; Bouizgarne 2013; Dey et al. 2014). Recent studies have focused on the culturable bacterial communities in corn, reporting that the most abundant bacteria linked to this plant belong to Firmicutes and *γ*-Proteobacteria, with the genera *Bacillus* and *Pseudomonas* the most frequently isolated microbes (Qaisrani et al. 2019). In addition, members of the Actinobacteria and Bacteroides have also been reported.

In a study on endophytes of healthy, asymptomatic corn plants, four strains with identification numbers JL103, JL109, PI101 and SS104 were isolated from corn tissues collected in Indiana, Iowa and Mississippi in the United States. Comparison of genome assemblies to strains within our culture collection isolated from other hosts showed that the novel species of JL109 and SS104 also occur on soybean and rice, respectively, in Arkansas. The type strains were characterized using molecular and phenotypic tests to determine their taxonomic placement. Our results indicate that the new strains represent four new species of the genera *Cellulomonas, Lelliottia, Paraburkholderia* and *Sphingomonas*, for which we propose the names *Cellulomonas zeae* sp. nov. (JL103)*, Lelliottia zeae* sp. nov. (SS104), *Paraburkholderia zeae* sp. nov. (PI101) and *Sphingomonas zeigerminis* sp. nov. (JL109).

## METHODS

### Isolation

Strains JL103, SS104 and JL109 were sourced from healthy field-grown *Zea mays* in Indiana, USA (JL103 and SS104) or Mississippi, USA (JL109). PI101 was isolated from ground, surface-sterilized seeds of an ancient land race *Zea mays*. Seeds were obtained from the North Central Regional Plant Introduction Station, Ames, Iowa. Plant tissue was washed with a mild detergent to remove particulates, surface-sterilized with bleach and ethanol, and homogenized. Serial dilutions of tissue homogenate were plated on a panel of media types for endophyte cultivation. Strain JL103, a small (0.7 mm diameter), pale yellow colony, arose on R2A after 5 days of incubation at 24°C. JL109, a bright yellow-pigmented, and PI101, a matte-white colony type, arose on Starch Casein agar after 7 and 3 days of incubation at 24°C, respectively. SS104 was isolated as a non-descript cream-colored colony on Potato Dextrose Agar. All strains were streaked to purity and stored in glycerol (20% v/v) at −80°C until subjected to further testing.

### Motility

The strains were tested for flagellar-dependent swimming and swarming motility on R2A plates solidified with 0.3% and 0.6% agar, respectively. Three independent colonies were inoculated onto R2A broth and grown for 36 hr at 24°C. Broth cultures were normalized to an OD600 of 0.1, and 1.5 μl of culture was spotted directly onto the surface of the motility agar. The diameter of colony expansion was measured for 5 days.

### Carbon source utilization

Substrate utilization was assessed using Biolog GenIII Microplates (Catalogue No. 1030) (Biolog Inc., Hayward, CA). Each bacterium was inoculated in duplicate plates using Protocol A, described by the manufacturer, with the exception that plates were incubated at 30°C. Respiration leading to reduction of the tetrazolium indicator was measured by absorbance at 590 nm.

### Biochemical analyses

Catalase activity was evaluated by immediate effervescence after the application of 3% (v/v) hydrogen peroxide solution via the tube method, a positive reaction was indicated by the production of bubbles. *Staphylococcus aureus* NCIMB 12702 and *Streptococcus pyogenes* ATCC 19615 were used as positive and negative controls, respectively. Oxidase activity was evaluated via the oxidation of Kovács oxidase reagent, 1% (w/v) tetra-methyl-p-phenylenediamine dihydrochloride in water, via the filter-paper spot method. A positive reaction was indicated when the microorganisms color changed to dark purple. *Pseudomonas aeruginosa* NCIMB 12469 and *Escherichia coli* ATCC 25922 were used as positive and negative controls, respectively. Urease activity was evaluated via the hydrolysis of urea in Christensen’s Urea Agar, using phenol red as a pH indicator. *Proteus hauseri* ATCC 13315 and *Escherichia coli* ATCC 25922 were used as positive and negative controls, respectively.

### Phylogenetic and genomic analyses

DNA was extracted from pure cultures using the Omega Mag-Bind Universal Pathogen Kit according to manufacturer’s protocol with a final elution volume of 60μl (Omega Biotek Inc., Norcross, GA). DNA samples were quantified using Qubit fluorometer (ThermoFisher Scientific, Waltham, MA) and normalized to 100 ng. DNA was prepped using Nextera DNA Flex Library Prep kit according to manufacturer’s instructions (Illumina Inc., San Diego, CA). DNA libraries were quantified via qPCR using KAPA Library Quantification kit (Roche Sequencing and Life Science, Wilmington, MA) and combined in equimolar concentrations into one 24-sample pool. Libraries were sequenced on a MiSeq using pair-end reads (2×200bp). Reads were trimmed of adapters and low-quality bases using Cutadapt (version 1.9.1) and assembled into contigs using MEGAHIT (version 1.1.2) (Li et al. 2015). Reads were mapped to contigs using Bowtie2 (version 2.3.4) (Langmead and Salzberg 2012), and contigs were assembled into scaffolds using BESST (2.2.8) (Sahlin et al. 2014).

16S rRNA gene sequences were extracted from genome assemblies using barrnap (Seemann 2019), and 16S rRNA gene phylogenetic analyses were performed using FastTree (Price et al. 2010) using a General Time Reversible substitution model. Taxon sampling for each species is described in the respective phylogenetic tree figure legend. GenBank accession numbers are provided in the phylogenetic tree figures.

Average nucleotide identity (ANI) analyses between genome assemblies were performed using the pyani ANIm algorithm (Richter and Rosselló-Móra 2009) implemented in the MUMmer package (Kurtz et al. 2004) retrieved from https://github.com/widdowquinn/pyani.

Geographic distribution and host range of novel species were inferred by ANI to assemblies from unidentified species from GenBank (Ciufo et al. 2018) and the Indigo internal collection. An ANI threshold of ≥95% indicated conspecificity (Chun et al. 2018; Richter and Rosselló-Móra 2009).

## RESULTS

### Phylogenetic and genomic analyses

#### *Cellulomonas zeae* sp. nov. strain JL103

Strain JL103 shared 99.3% 16S rRNA gene sequence identity with *Cellulomonas humilata* ATCC 25174^T^, followed by *Cellulomonas xylanilytica* XIL 11^T^ (99.2%) and shared less than 98% sequence similarity with the remaining *Cellulomonas* species. A phylogenetic tree using FastTree (Price et al. 2010) confirmed the affiliation of strain JL103 to the genus *Cellulomonas*. JL103 formed a monophyletic group with the species *C. humilata* and *C. xylanilytica* supported by high bootstrap support (Figure 1) and showing that it could be distinguished from its phylogenetic neighbors. Average nucleotide identity (ANI) values between *C. humilata* ATCC 25174^T^, *C. xylanilytica* NBRC 101102^T^ and the new isolate was 86.7 and 86.5%, respectively. Their values are well below the threshold for species demarcation (Richter and Rosselló-Móra 2009; Chun et al. 2018) providing further genomic support that strain JL103 represents a new genomic species of *Cellulomonas*.

**Figure 1.**
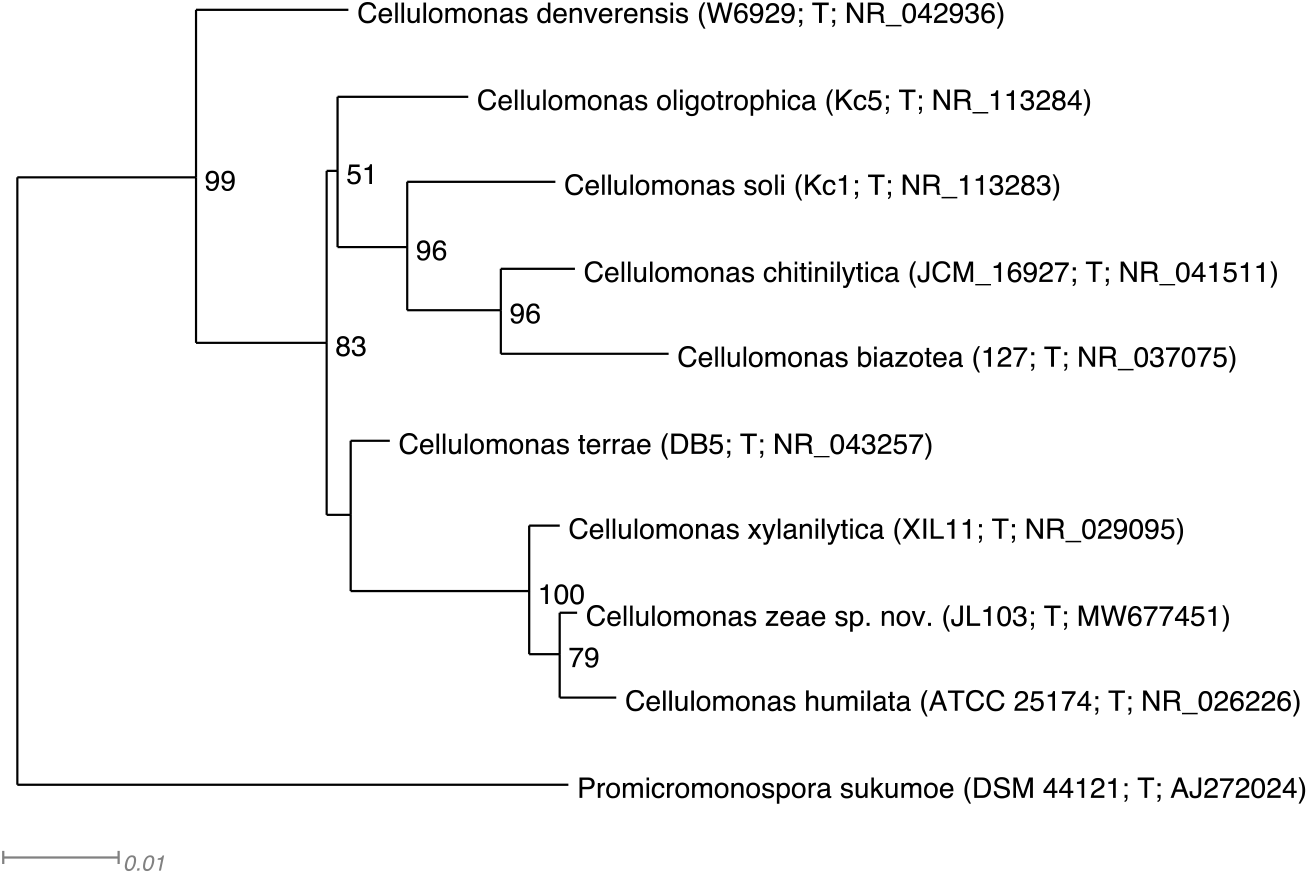
Phylogenetic 16S tree of *Cellulomonas zeae* sp. nov. and relatives generated using FastTree (Price et al. 2010). The eight valid species with the highest 16S identities to *C. zeae* were included in the tree, the tree is rooted with *Promicromonospora sukumoe*. Strain numbers and GenBank accession numbers follow species names, T stands for ‘type’. Support values above 50% are given by the branches. *Cellulomonas zeae* is most closely related to *C. humilata* and *C. xylanilytica* with 100% support. Branch lengths are proportional to the changes along the branches, a scale bar is provided at the bottom.

#### *Lelliottia zeae* sp. nov. strain SS104

16S rRNA sequence identity between strain SS104 and the type strains *Lelliottia amnigena* JCM1237^T^, *Lelliottia jeotgali* PFL01^T^ and *Lelliottia nimipressuralis* LMG 10245^T^ were 99.5%, 98.9% and 98.3%, respectively. The phylogenetic position of SS104 suggested allocation to the genus *Lelliottia* (Figure 2). Strain SS104 formed a monophyletic group with *Lelliottia amnigena*, supported by bootstrap support of 88% (Figure 2). The two top ANI hits were to genomes of *Lelliottia amnigena* and *Lelliottia jeotgali* with 93.7% and 86.0%, respectively. These results indicate that strain SS104 represents a new genomic species of *Lelliottia*.

**Figure 2.**
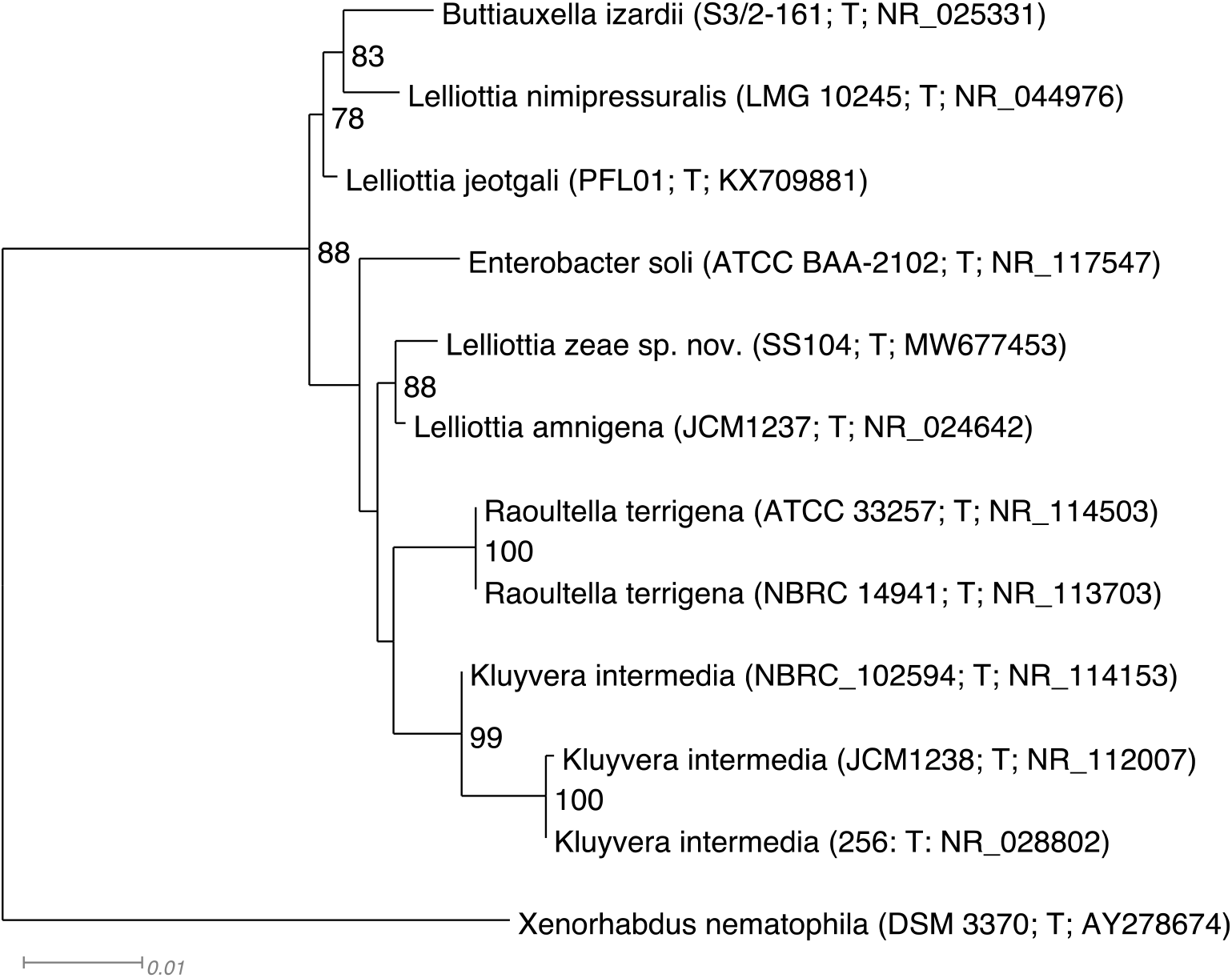
Phylogenetic 16S tree of *Lelliottia zeae* sp. nov. and relatives generated using FastTree (Price et al. 2010). The 10 valid s with the highest 16S identities to *L. zeae* were included in the tree, the tree is rooted with *Xenorhabdus nematophila*. Strain numbers and GenBank accession numbers follow species names, T stands for ‘type’. Support values above 70% are given by the branches. *Lelliottia zeae* is most closely related to *Lelliottia amnigena* with 88% support. Branch lengths are proportional to the changes along the branches, a scale bar is provided at the bottom.

#### *Paraburkholderiae zeae* sp. nov. strain PI101

16S rRNA sequence identity between strain PI101and the type strains *Paraburkholderia strydomiana* WK1f ^T^ and *Paraburkholderia aromaticivorans* BN5 ^T^ were 98.5% and 98.3%, respectively. The phylogenetic position of SS104 suggested allocation to the genus *Paraburkholderia* (Figure 3). Strain PI101formed a monophyletic group with the type strain *Paraburkholderia sediminicola* HU2-65W ^T^, supported by bootstrap support of 91% (Figure 3). The two top ANI hits were to the genome of *Paraburkholderia aromaticivorans* and the genome of the type strain *Paraburkholderia xenovorans* LB400 ^T^ with 91.5% and 90.8%, respectively. These results indicate that strain PI101 represents a new genomic species of *Paraburkholderia*.

**Figure 3.**
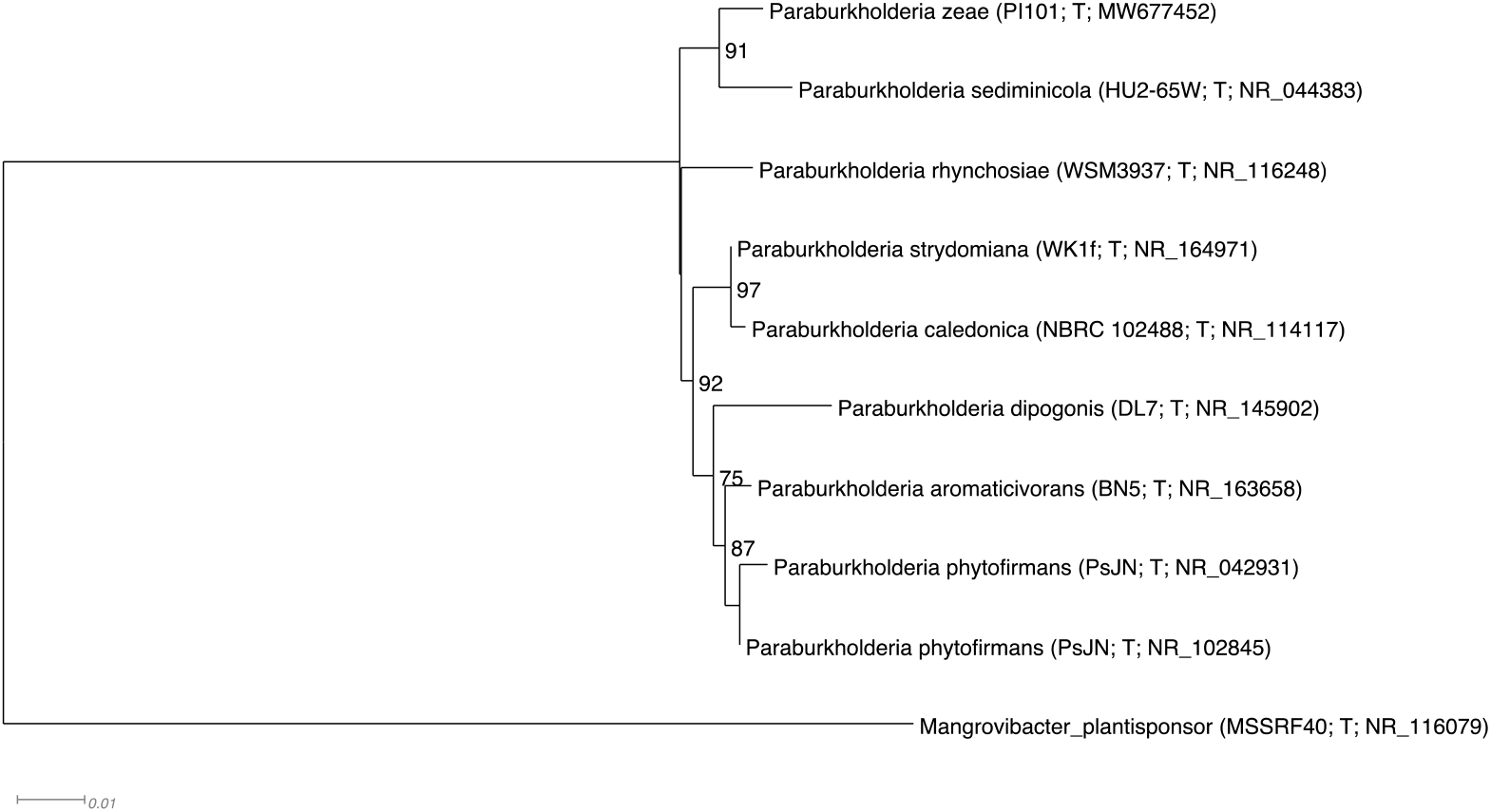
Phylogenetic 16S tree of *Paraburkholderia zeae* sp. nov. and relatives generated using FastTree (Price et al. 2010). The 8 valid entries with the highest 16S identities to *P. zeae* were included in the tree, the tree is rooted with *Mangrovibacter plantisponsor*. Strain numbers and GenBank accession numbers follow species names, T stands for ‘type’. Support values above 70% are given by the branches. *Paraburkholderia zeae* is most closely related to *Paraburkholderia sediminicola* with 91% support. Branch lengths are proportional to the changes along the branches, a scale bar is provided at the bottom.

#### *Sphingomonas zeigerminis* sp. nov. strain JL109

The strain JL109 16S rRNA was 100% identical to *Sphingomonas trueperi* NBRC 100456^T^. The phylogenetic position of JL109 confirmed its allocation in the genus *Sphingomonas* (Figure 4) and was most closely related to *S. trueperi*, *Sphingomonas azotifigens* Y39^T^ and *Sphingomonas pituitosa* DSM 13101^T^ (Figure 4). ANI values between the genome sequence of isolate JL109 and the related species *S. azotifigens, S. pituitosa* and *S. trueperi* were 89.8%, 89.4% and 92.2%, respectively. These results indicate that strain JL109 represents a new genomic species of *Sphingomonas*.

**Figure 4.**
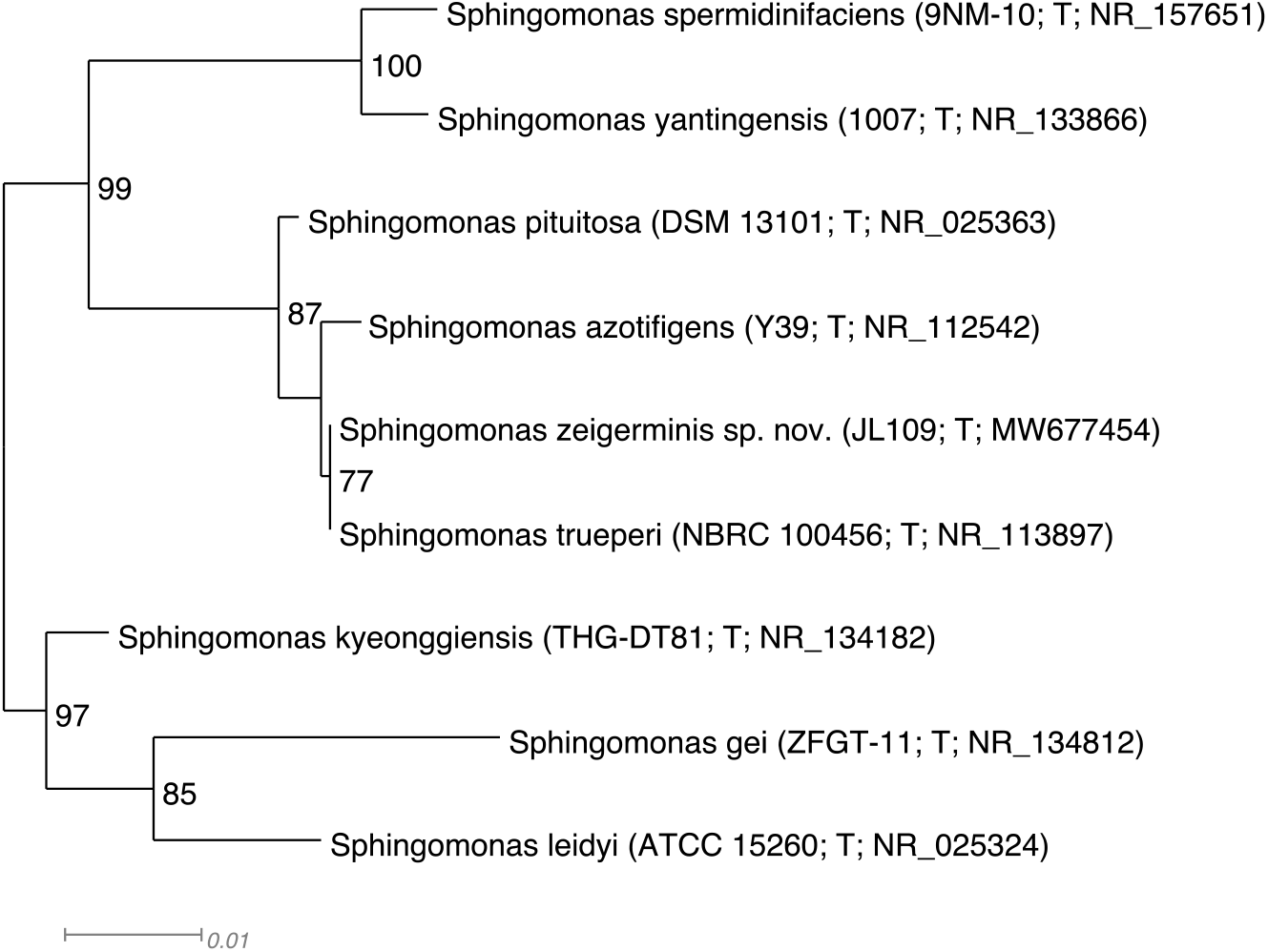
Phylogenetic 16S tree of *Sphingomonas zeigerminis* sp. nov. and relatives generated using FastTree (Price et al. 2010). The eight valid species with the highest 16S identities to *S. zeigerminis* were included in the tree that was midpoint-rooted. Strain numbers and GenBank accession numbers follow species names, T stands for ‘type’. Support values above 70% are given by the branches. *Sphingomonas zeigerminis* is most closely related to *S. trueperi, S. azotifigens* and *S. pituitosa* with 87% support. Branch lengths are proportional to the changes along the branches, a scale bar is provided at the bottom.

### Geographic distribution and host range

Geographic distribution and host range of the novel species was inferred by comparison to congeneric genome assemblies of unidentified species from GenBank and the Indigo internal collection. Hits to respective novel species included the Indigo strains *Lelliottia zeae* strain SS115 (ANI: 98.5%; query coverage: 94.6%) collected from healthy *Oryza sativa* plants in Arkansas, and *Sphingomonas zeigerminis* strain VK114 (95.8%; 81.4%) from healthy *Glycine max* in Arkansas. No hits were found among GenBank assemblies from environmental and clinical sources. Known geographic distributions of the novel species from culturing is illustrated in Figures 5, 6, 7 and 8, and host range in Table 1.

**Figure 5.**
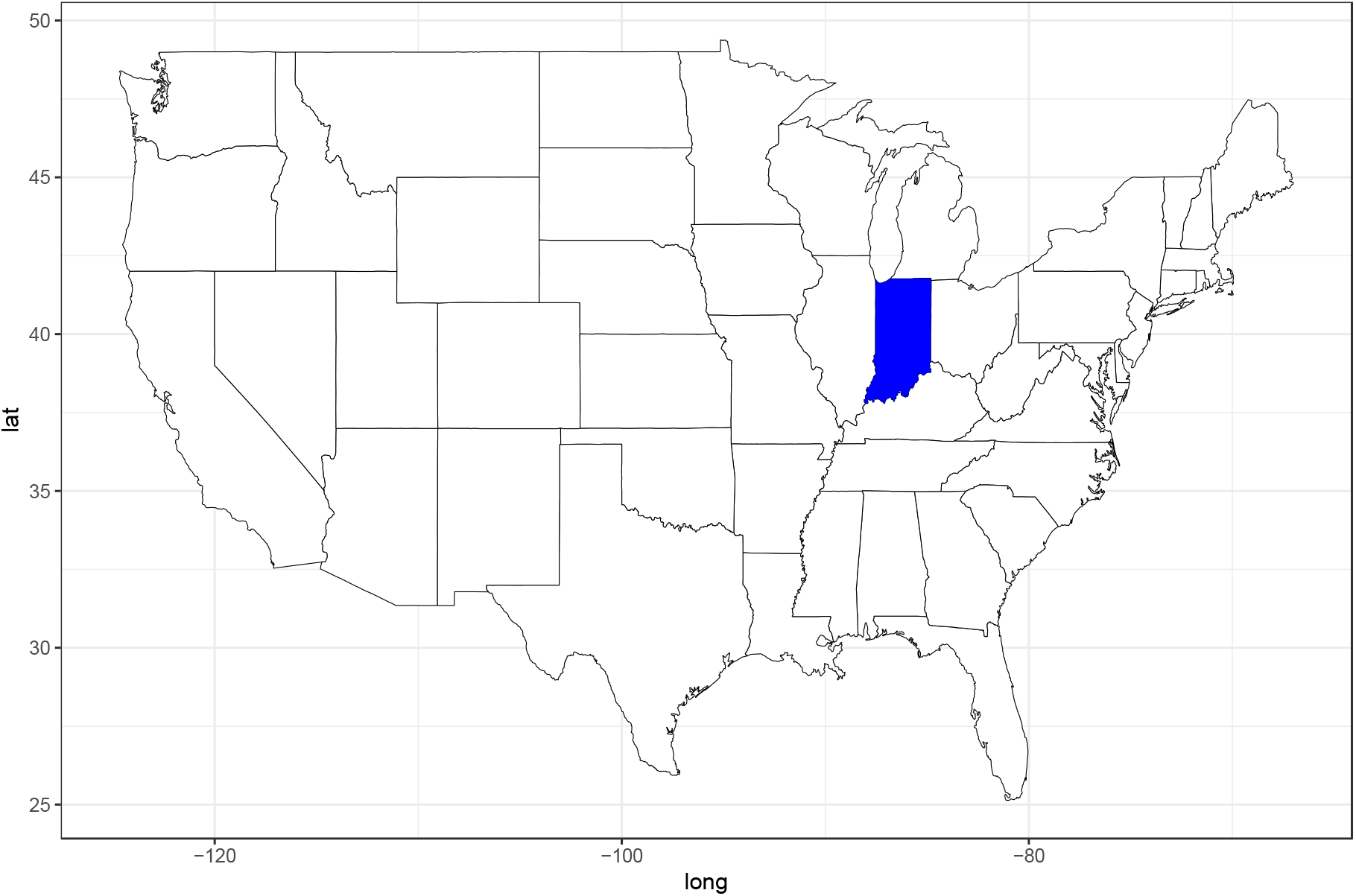
Known geographic distribution of *Cellulomonas zeae* sp. nov. based on culturing. Dark blue marks state where type strain was collected, no cultures are known from other states.

**Figure 6.**
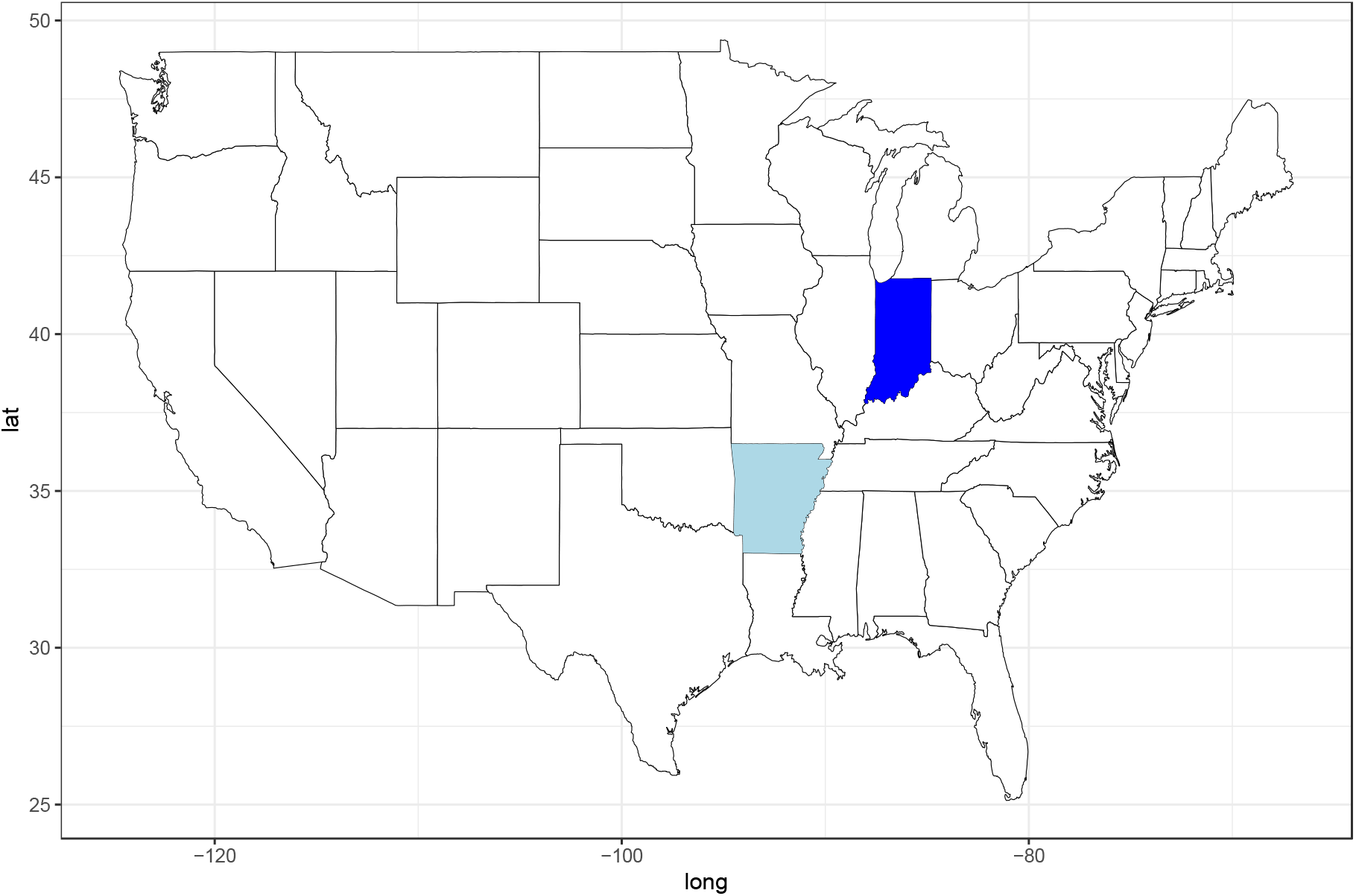
Geographic distribution of *Lelliottia zeae* sp. nov. based on culturing. Colors mark states where species was collected. Dark blue indicates type strain, light blue additional strains.

**Figure 7.**
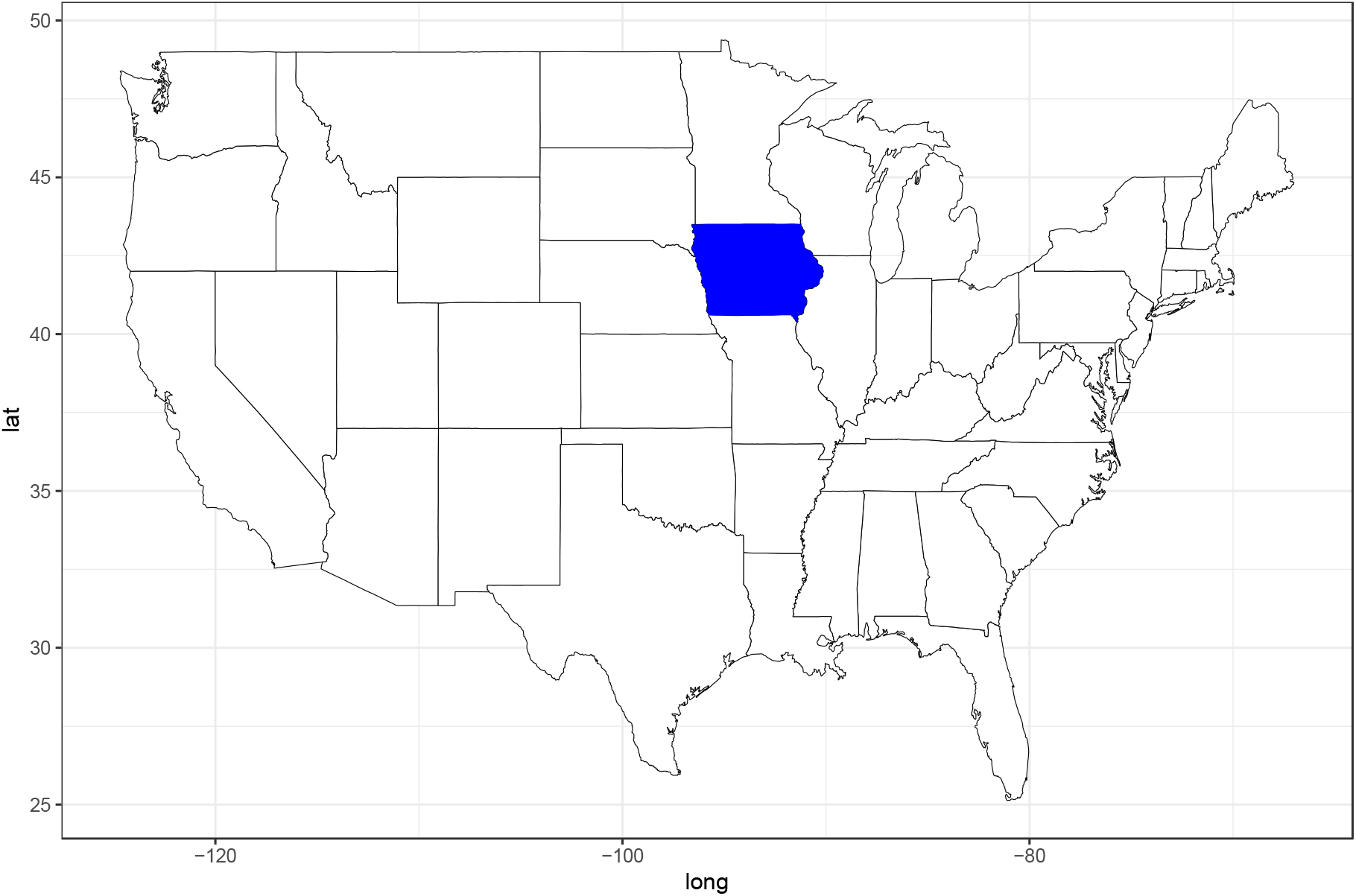
Geographic distribution of *Paraburkholderia zeae* sp. nov. based on culturing. Dark blue marks state where type strain was collected, no cultures are known from other states.

**Figure 8.**
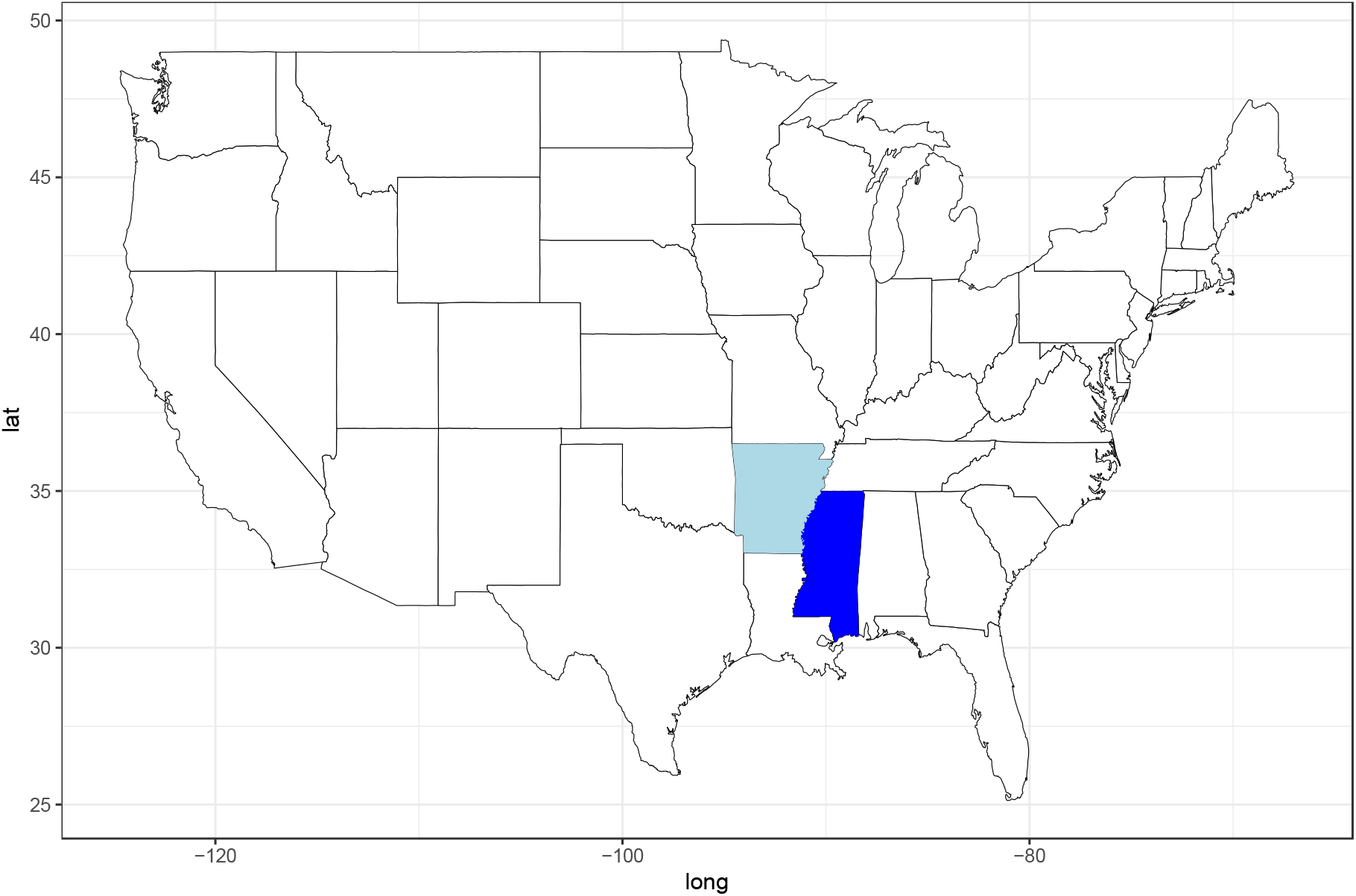
Geographic distribution of *Sphingomonas zeigerminis* sp. nov. based on culturing. Colors mark states where species was collected. Dark blue indicates type strain, light blue additional strains.

**Table 1.** Hosts of novel species based on culturing.

| **Species** | **Host<sup>1</sup>** |
| --- | --- |
| *Cellulomonas zeae* | *Zea mays* |
| *Lelliottia zeae* | *Oryza sativa*, *Zea mays* |
| *Paraburkholderia zeae* | *Zea mays* |
| *Sphingomonas zeigerminis* | *Glycine max*, *Zea mays* |

<sup>1</sup> Strains originated from asymptomatic plant tissues.

### Morphology, physiology and biochemical characteristics

#### *Cellulomonas zeae* sp. nov. strain JL103

Cells of strain JL103 stained Gram-positive and were non-spore forming rod-shaped with an average cell size of 0.4 μm wide and 1.3-1.5 μm long. They grew as single cells or in pairs (Figure 9) but no motility or swarming was observed. These characteristics are consistent with the description of the genus *Cellulomonas* (Stackebrandt and Schumann 2012). Good growth was observed on nutrient agar (NA), tryptic soy agar (TSA) and R2A agars at 30°C after 7 days. Abundant growth was observed on R2A after 48 hrs. On this medium the strain also produced an exopolysaccharide. Colonies were pale yellow, circular and entire. Except for R2A agar, growth at 22°C was weak on these media. The strain is also able to grow in the presence of 4% NaCl, unlike *C. humilata* ATCC 25174^T^. Other differential characteristics between JL103 and the species *C. humilata* and *C. xylanilytica* are found in Table 2. Additional physiological characteristics are found in the species description.

**Figure 9.**
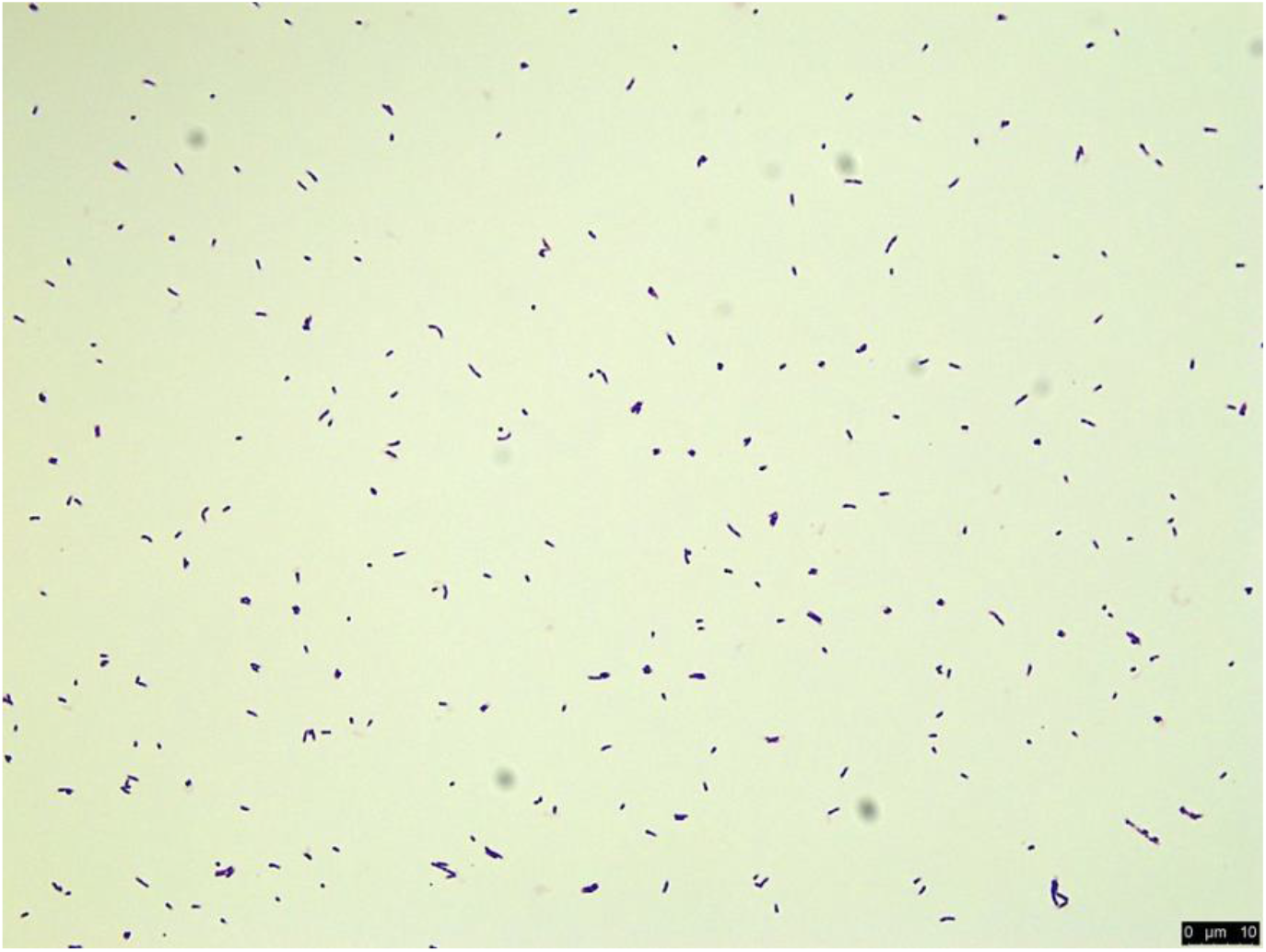
Morphology of *Cellulomonas zeae* sp. nov. JL103 depicted following Gram stain under bright field microscopy. Bar = 10 μm

**Table 2.** Physiological characteristics of *Cellulomonas zeae* sp. nov. strain JL103 and related *Cellulomonas* species.

|  | *C. zeae* J1103<sup>T</sup> | *C. xylanilytica*<br>XIL 11<sup>T</sup> | *C. humilata*<br>ATCC 25174<sup>T</sup> |
| --- | --- | --- | --- |
| **General characteristics:** |  |  |  |
| **Colony morphology** | Pale yellow, smooth, no mycelium | Lemon yellow, smooth, no mycelium | White, with branching mycelium |
| **Growth at 30°C** | + | + | + |
| **Growth at 37°C** | - | + | Poor to no growth |
| **Growth with 4% NaCl** | + | nr | - |
| **Motility** | - | - | - |
| **Oxidase** | - | + | - |
| **Catalase** | - | + | - |
| **Utilization of:** |  |  |  |
| **Acetate** | - | - | - |
| **Gluconate** | - | - | + |
| **D-Lactose** | - | + | + |
| **Lactate** | - | - | - |
| **D-Mannose** | + | + | + |
| **D-Mannitol** | - | - | + |
| **L-Rhamnose** | w | + | + |
| **Gentiobiose** | - | + | nr |
| **Maltose** | + | + | nr |
| **N-Acetyl-D-Glucosamine** | - | + | nr |

+, Positive; w, Weakly positive; -, Negative; nr, Not reported

#### *Lelliottia zeae* sp. nov. strain SS104

Cells stained Gram-negative, were rod-shaped and measured 0.8-1 μm wide and 2 μm long (Figure 10). They were motile by swimming and did not produce spores. Colonies had a cream color and were circular, smooth and raised. Good growth was observed after 24 h on TSA, NA and R2A, at 22 and 30°C. Similar to other *Lelliottia* especies, the new strain is catalase negative and oxidase positive, but unlike its closest phylogenetic neighbor, *L. amnigena, L. zeae* SS104 does not produce urease. Temperature is also useful to differentiate between *L. zeae* and the species *L. jeotgali* and *L. amnigena*. The new strain grows at 41°C but not at 45°C, while *L. jeotgali* grows at both temperatures and *L. amnigena* does not show any growth at these temperatures. Additional differential characteristics based on carbon source assimilation are found in Table 3.

**Figure 10.**
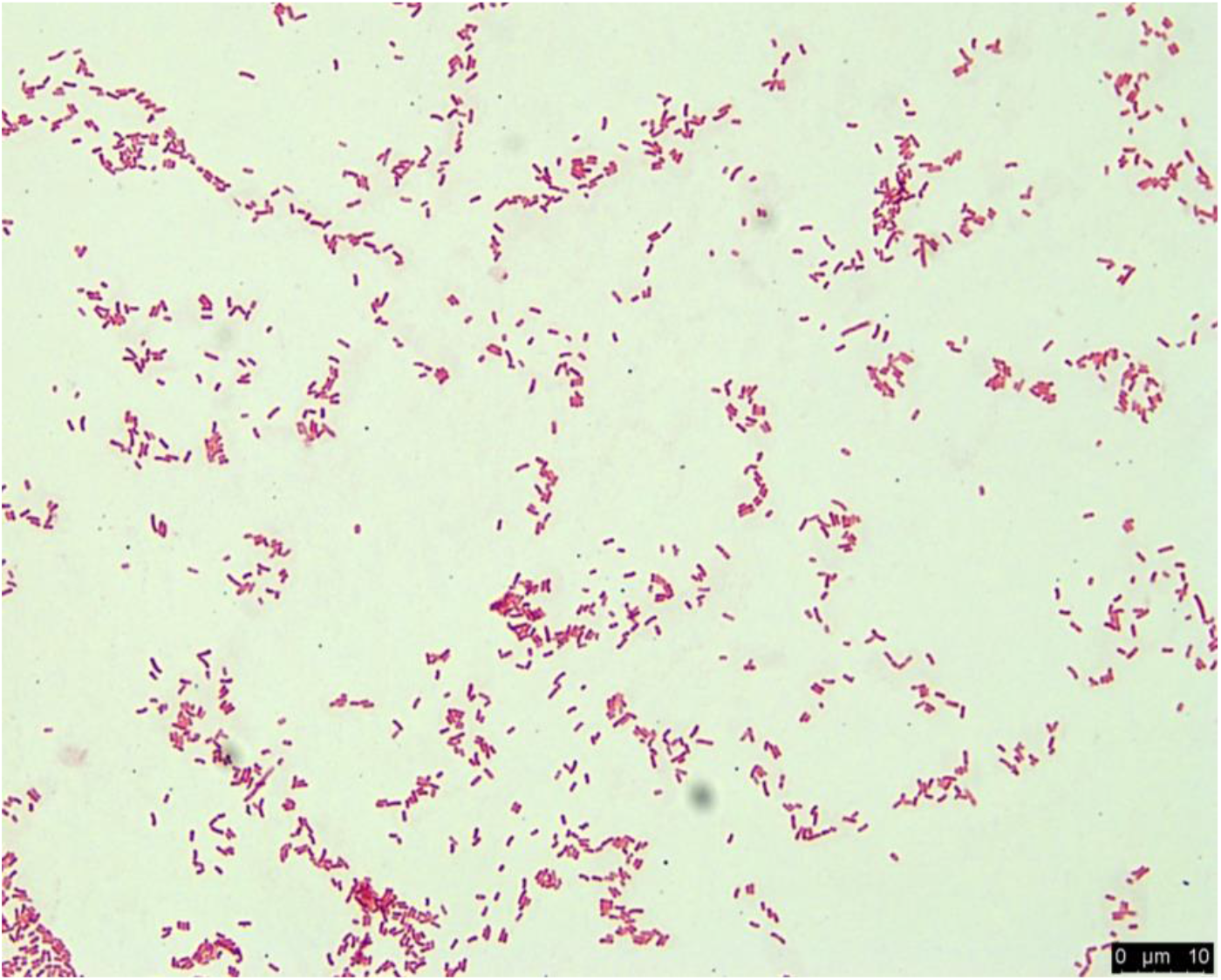
Morphology of *Lelliottia zeae* sp. nov. strain SS104 depicted following Gram stain using bright field microscopy. Bar = 10 μm.

**Table 3.**
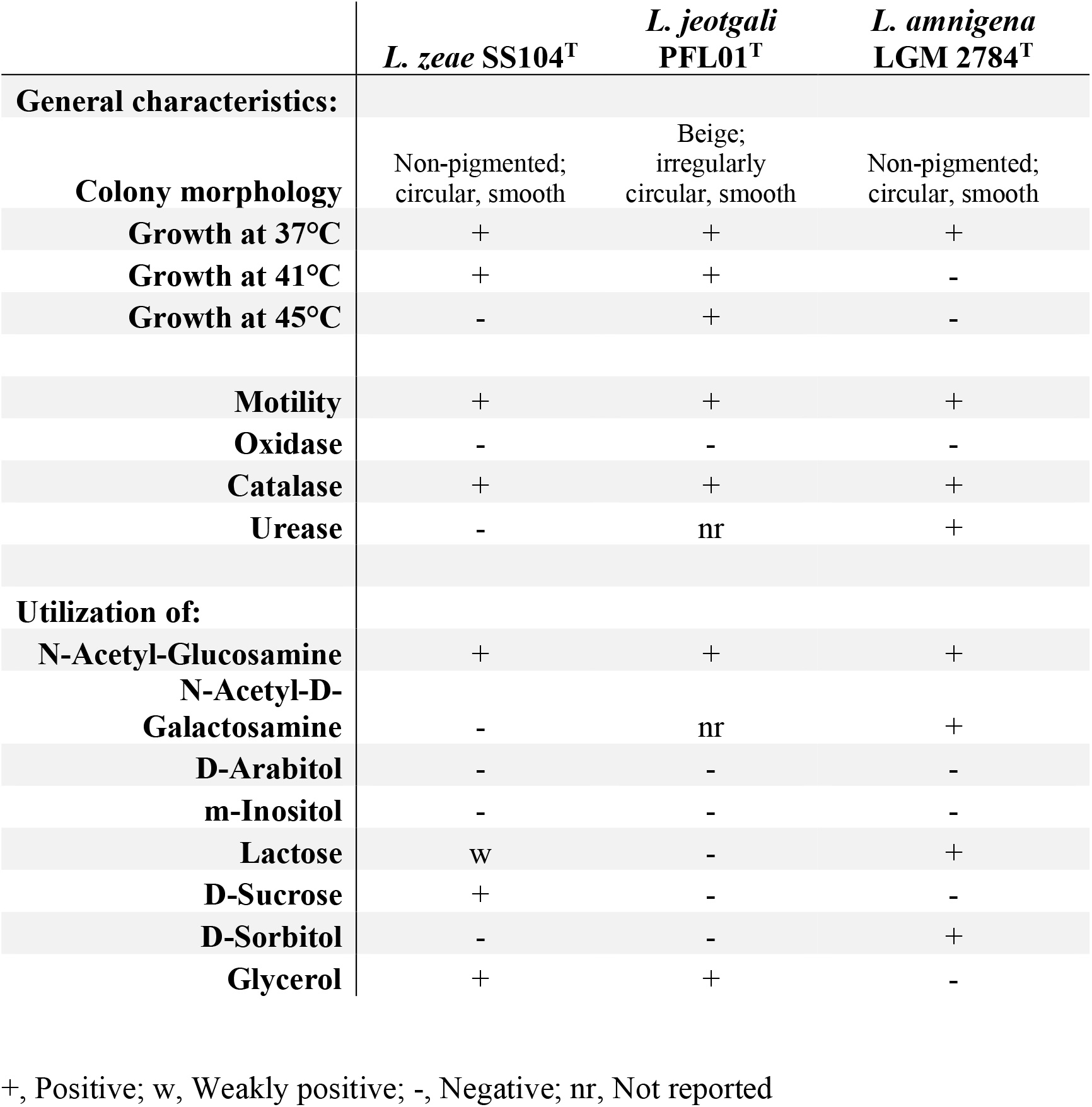
Physiological characteristics of *L. zeae* and related *Lelliottia* species.

|  | *L. zae* SS104<sup>T</sup> | *L. jeotgali*<br>PFL01<sup>T</sup> | *L. amnigena*<br>LGM 2784<sup>T</sup> |
| --- | --- | --- | --- |
| **General characteristics:** |  |  |  |
| **Colony morphology** | Non-pigmented;<br>circular, smooth | Beige;<br>irregularly<br>circular, smooth | Non-pigmented;<br>circular, smooth |
| **Growth at 37°C** | + | + | + |
| **Growth at 41°C** | + | + | - |
| **Growth at 45°C** | - | + | - |
| **Motility** | + | + | + |
| **Oxidase** | - | - | - |
| **Catalase** | + | + | + |
| **Urease** | - | nr | + |
| **Utilization of:** |  |  |  |
| **N-Acetyl-Glucosamine** | + | + | + |
| **N-Acetyl-D-Galactosamine** | - | nr | + |
| **D-Arabitol** | - | - | - |
| **m-Inositol** | - | - | - |
| **Lactose** | w | - | + |
| **D-Sucrose** | + | - | - |
| **D-Sorbitol** | - | - | + |
| **Glycerol** | + | + | - |

+, Positive; w, Weakly positive; -, Negative; nr, Not reported

#### *Paraburkholderia zeae* sp. nov. strain PI101

Cells of PI101 are Gram stain-negative, straight or slightly curved non-spore forming rods. Cell size ranges from 0.5-1.2 μm in width and 1.2 to 3.0 μm in length (Figure 11). Motility, which is characteristic of *Paraburkholderia* species, was observed on 0.3% agar. Good growth at 22 and 30°C is observed after 24 h on NA, TSA and R2A agars, with exopolysaccharide production on the latter medium. On these media, colonies were white, round, and smooth. The new strain produced catalase and oxidase; these enzymes are also produced by its phylogenetic neighbors, *P. aromaticivorans, P. strydomiana and P caledonica*, except that *P. caledonica* is negative for oxidase. Interestingly, only *P. zeae*, PI101, produced urease. Other physiological characteristics, including assimilation of several carbon sources and comparison with its phylogenetic neighbors are found in Table 4.

**Figure 11.**
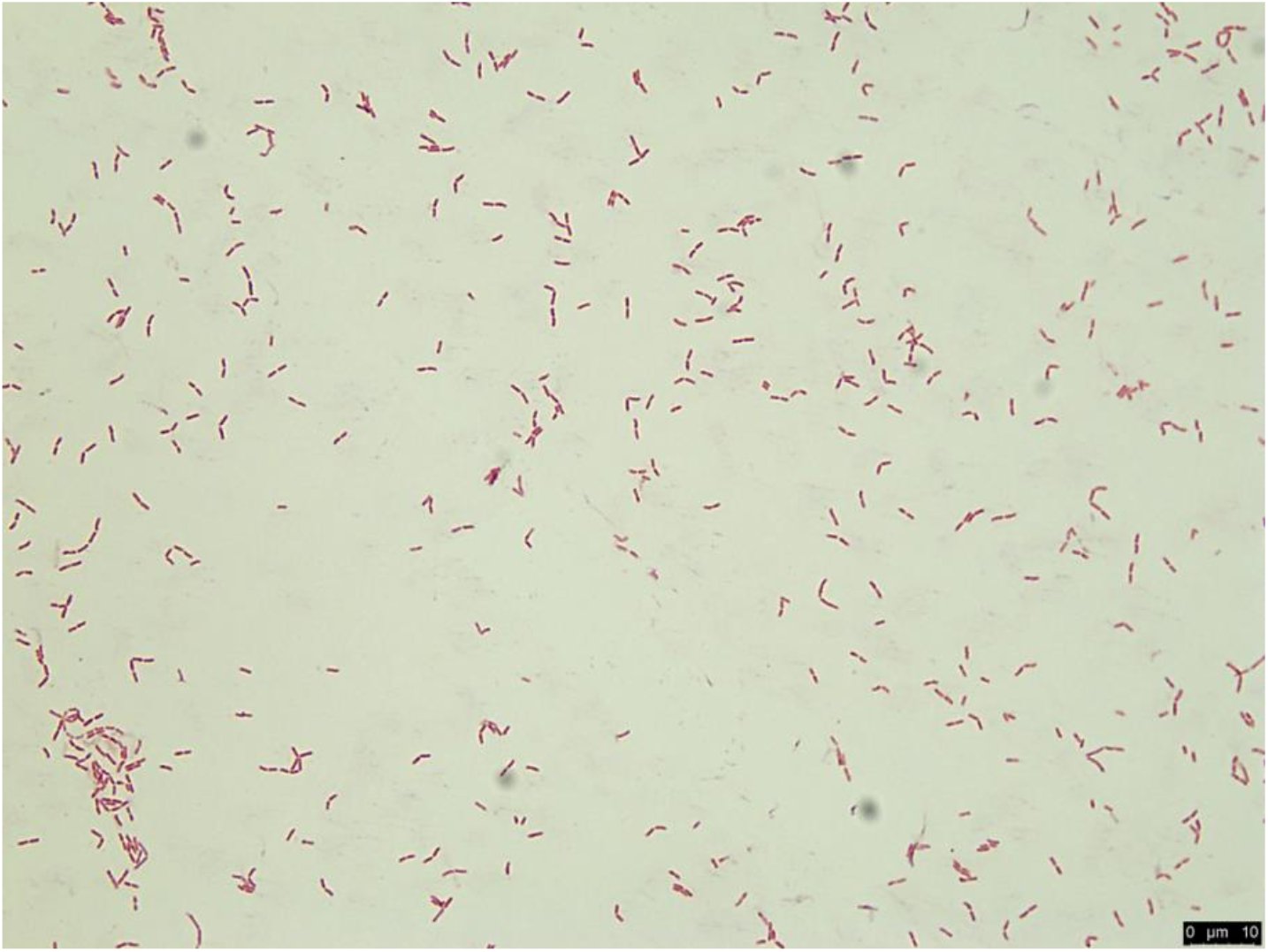
Morphology of *Paraburkholderia zeae* sp. nov. strain PI101 depicted following Gram stain using bright field microscopy. Bar = 10 μm.

**Table 4.** Physiological characteristics of *Paraburkholderia zeae* and related *Paraburkholderia* species.

|  | *P.*<br>*zeae* | *P.*<br>*aromaticiv.* | *P.*<br>*strydomia.* | *P.*<br>*caledonica* |
| --- | --- | --- | --- | --- |
| **General characteristics:** |  |  |  |  |
| **Colony morphology** | Non-pigmented;<br>round, smooth | Non-pigmented;<br>round, smooth | Non-pigmented;<br>round, smooth | nr |
| **Growth at 37°C** | + | + | + | - |
| **Oxidase** | + | + | + | - |
| **Catalase** | + | + | + | + |
| **Urease** | + | - | - | - |
| **Utilization of:** |  |  |  |  |
| **Sucrose** | - | + | - | - |
| **Cellobiose** | - | + | + | nr |
| **Melibiose** | - | + | nr | nr |
| **Raffinose** | - | + | - | nr |
| **D-Glucose** | + | + | + | + |
| **Lactose** | - | + | - | + |
| **N-Acetyl-D-Galactosamine** | - | + | - | + |
| **N-Acetyl-D-Glucosamine** | + | + | + | + |
| **m-inositol** | + | + | - | + |
| **D-Mannitol** | + | + | + | + |
| **D-Sorbitol** | + | + | + | nr |
| **Citric acid** | w | + | - | - |
| **Propionic acid** | - | + | + | nr |
| **Acetic acid** | - | nr | + | nr |

+, Positive; w, Weakly positive; -, Negative; nr, Not reported
Taxa: *P. zeae*: *P. zeae* PI101<sup>T</sup>; *P. aromaticiv.*: *P. aromaticivorans* BN5<sup>T</sup>; *P. strydomia.*: *P. strydomiana* WK1.1f<sup>T</sup>; *P. cal.*: *P. caledonica* LMG19076<sup>T</sup>

#### *Sphingomonas zeigerminis* sp. nov. strain JL109

Cells of strain JL109 stained Gram-negative, and are slightly curved or ovoid rod shape. Cell size ranges from 0.3-1.2 μm x 0.5 −3.0 μm (Figure 12). Colonies were wrinkled, entire, raised and bright yellow on Nutrient Agar. The new strain was motile by flagella and strictly aerobic, with an oxidative metabolism. Growth was good between 30-37 °C, weak in the presence of 1% (w/v) NaCl and did not grow below pH 6. The new strain produced oxidase as many other *Sphingomonas* species. Catalase production and use of acetate can be used to differentiate strain JL109 from *S. azotifigens* Y39^T^, *S. pituitosa* DSM 13101^T^ and *S. trueperi* NBRC 100456^T^. Table 5 presents a list of additional characteristics that can be useful to differentiate strain JL109 from its closest phylogenetic neighbors. In addition, a detailed description of other physiological characteristics is given in the species protologue.

**Figure 12.**
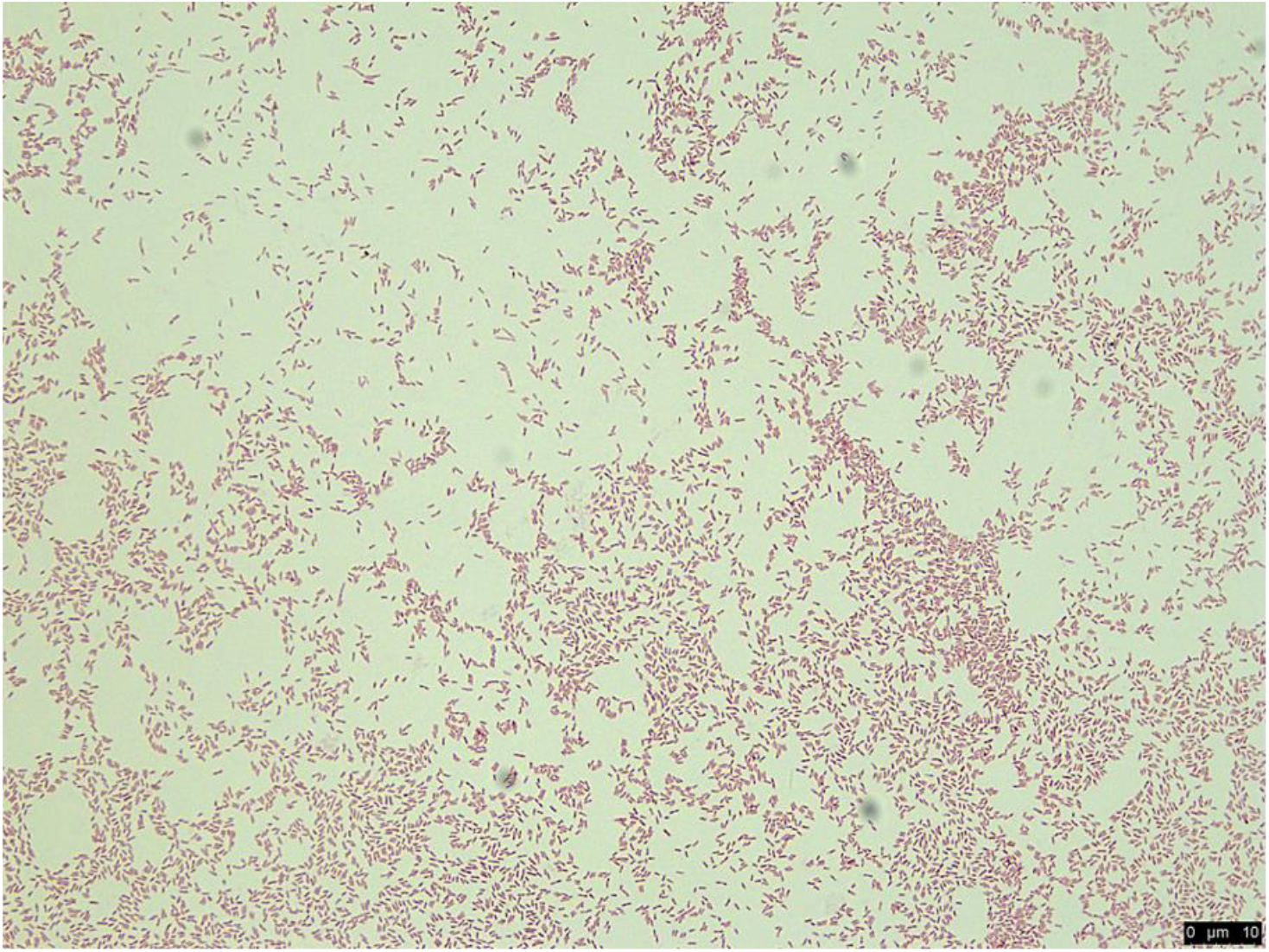
Morphology of *Sphingomonas zeigerminis* JL109 depicted following Gram stain and under bright field microscopy. Bar = 10 μm.

**Table 5.**
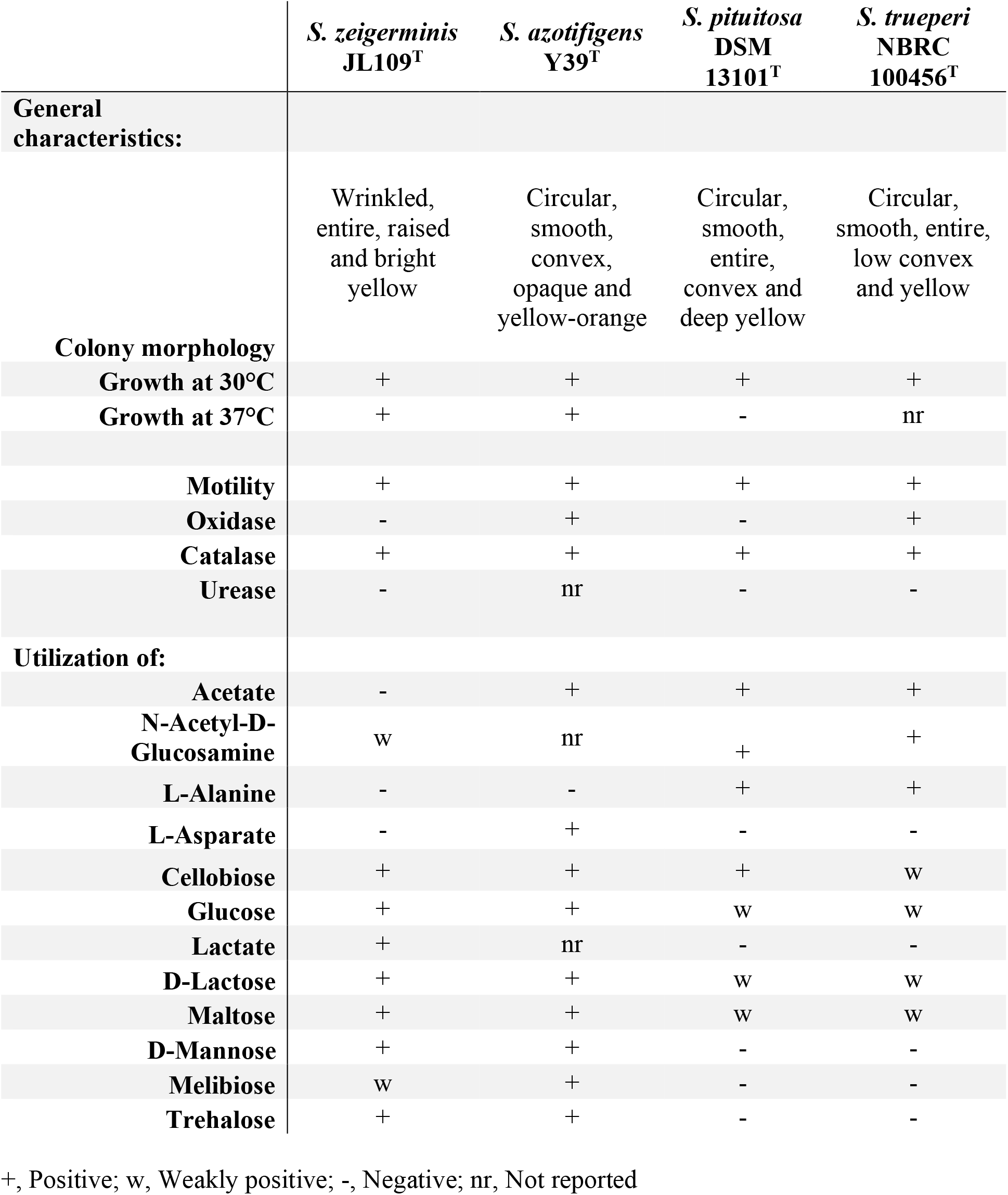
Physiological characteristics of *Sphingomonas zeigerminis* sp. nov. strain JL109 and related *Sphingomonas* species.

## DESCRIPTION OF CELLULOMONAS ZEAE SP. NOV

*Cellulomonas zeae* sp. nov. *zeae*, ze’ae. L fem. n. zeae of *Zea* (maize), isolated from *Zea mays*

Cells are Gram-stain positive, non-flagellated, non-sporeforming, coccoid or rod-shaped cells (1.3-1.5 μm x 0.4 μm). growth is aerobic or facultatively anaerobic and chemo-organotrophic. Colonies on R2A and NA are pale yellow, smooth, circular, entire. Good growth on R2A agar after 48 hrs and 30°C. EPS is produced on R2A. Aerial mycelium is not produced. Cells grow between 24-37°C (optimum 30°C); grows at pH 6-7 but not at pH 5. *C. zeae* grows in the presence of 0-7% (w/v) NaCl and is catalase and oxidase negative. Substrates used as carbon sources include dextrin, maltose, D-trehalose, D-cellobiose, D-turanose, stachyose, D-raffinose, D-melibiose, D-salicin, α-D-glucose, D-mannose, D-fructose, D-galactose, L-rhamnose, L-acetoacetic acid.

The type strain is resistant to the following antibiotics: troleandomycin, rifamycin sv, minocycline, lincomycin, vancomycin, nalidixic acid and aztreonam.

The type strain, JL103^T^, was isolated from corn roots collected in Indiana, USA.

## DESCRIPTION OF LELLIOTTIA ZEAE SP. NOV

*Lelliottia zeae* sp. nov. *zeae*, ze’ae. L fem. n. zeae of *Zea* (maize), isolated from *Zea mays*

This strain is a Gram-stain negative, non-sporeforming, motile rod (0.8-1 x 2 μm). Growth is aerobic with an oxidative metabolism. Colonies on R2A agar are cream color. Growth is observed at pH 5 and 6 and in the presence of 1-4% NaCl (w/v); weak growth is observed at 8% NaCl. The following substrates are used as carbon sources: D-maltose, D-trehalose, D-cellobiose, D-gentobiose, sucrose, D-raffinose, D-melibiose, β-Methyl-D-glucoside, D-salicin, N-Acetyl-D-glucosamine, N-Acetyl-β-D-mannosamine, N-Acetyl-Neuraminic acid, α-D-glucose, D-mannose, D-fructose, D-galactose, L-rhamnose, inosine, D-mannitol, glycerol, D-glucose-PO4, D-fructose-PO4, proline, arginine, L-glutamic acid, L-serine, pectin, D-galacturonic acid, citric acid, malic acid.

The type strain is resistant to the following antibiotics: troleandomycin, rifamycin sv, lincomycin, vancomycin, and aztreoman but susceptible to minocycline and nalidixic acid.

The type strain, SS104^T^, was isolated from corn tissue in Indiana, USA.

## DESCRIPTION OF PARABURKHOLDERIA ZEAE SP. NOV

*Paraburkholderia zeae* sp. nov. *zeae*, ze’ae. L fem. n. zeae of *Zea* (maize), isolated from *Zea mays*

Cells are Gram-negative, motile, non-sporulating, rods (0.8 μm by 1.4 μm). The strain grows strictly aerobic using an oxidative metabolism. Colonies are white, round, and smooth. Growth is observed at pH 5 and 6. *P. zeae* grows in the presence of 1% (w/v) NaCl, but not at 4 or 8%. The following substrates are used as carbon sources: N-acetyl-D-glucosamine, D-glucose, D-mannose, D-fructose, D-galactose, D-fucose, L-fucose, L-rhamnose, D-sorbitol, D-mannitol, D-arabitol, *myo*-inositol, glycerol, D-glucose-PO4, D-fructose-PO4, glycyl-L-proline, L-alanine, L-arginine, L-glutamic acid, L-histidine, L-serine, D-gluconic acid, D-glucuronic acid, quinic acid, p-hydroxy-phynylacetic acid, methyl pyruvate, L-lactic acid, L-malic acid, γ-amino-butyric acid, β-hydroxyl-D,L-butyric acid.

The type strain is resistant to rifamycin sv, lincomycin, vancomycin, and aztreoman, but susceptible to minocycline, and nalidixic acid.

The type strain, PI101^T^ was isolated from corn seeds, Iowa, USA.

## DESCRIPTION OF SPHINGOMONAS ZEIGERMINIS SP. NOV

*Sphingomonas zeigerminis* (ze.i.ger’mi.nis. L. fem. n. zea spelt and the generic name of corn (*Zea*); L. neut. n. germen a sprout; N.L. gen. n. zeigerminis of a corn seedling).

Cells are Gram-stain negative, motile, non-spore forming rods. Slightly curved or ovoid rods. Bright yellow, wrinkled, entire and raised colonies. Good growth is obtained at 30°C and pH 7. Weak growth is observed on 1% NaCl. The following substrates are used as carbon sources: dextrin, D-maltose, D-trehalose, D-cellobiose, gentobiose, sucrose, α-lactose, α-D-glucose, D-mannose, D-galactose, sodium lactate, L-galactonic acid lactone, D-glucuronic acid and L-malic acid. Weak growth on turanose, D-melibiose, N-acetyl-D-glucosamine, N-acetyl-D-galactosamine, DL-fucose, pectin, D-galacturonic acid, L-malic acid and Tween 40. No growth on stachyose, D-raffinose, D-salicin, D-fructose, 3-methyl-glucose, L-rhamnose, inosine, D-serine, D-sorbitol, D-mannitol, D-arabitol, *myo*-inositol, glycerol, D-aspartic acid, D-serine. Gelatin is not hydrolyzed, but pectin is weakly degraded.

The type strain is resistant to lincomycin, troleandomycin, rifamycin SV, nalidixic acid and aztreonam. Sensitive to vancomycin.

The type strain, JL109^T^, was isolated from corn tissue in Mississippi, USA.

## ACKNOWLEDGEMENTS

We would like to thank Professor Aharon Oren, The Hebrew University of Jerusalem, for help with nomenclature.

## Notes

### Competing Interest Statement

The authors have declared no competing interest.

### Summary of Updates

Geographic distribution and host range of the new species was updated based on genome comparison to other strains from culture. And erroneous numbering of figure references in text was corrected.

